# A framework for automated multimodal HDX-MS analysis

**DOI:** 10.1101/2025.03.13.643099

**Authors:** Lisa M. Tuttle, Rachel E. Klevit, Miklos Guttman

## Abstract

We present pyHXExpress, a customizable codebase for automated high-throughput multimodal analysis of all spectra generated from HDX-MS experiments. The workflow was validated against a synthetic test dataset to test the fitting algorithms and to confirm the statistical outputs. We further establish a framework for the determination of multimodality throughout a protein system by rigorous evaluation of multimodal fits across all peptide spectra. We demonstrate this approach using entire protein datasets to detect multimodality, conformational heterogeneity, and characterize dynamics of small heat shock protein HSPB5 and two disease mutants.

## INTRODUCTION

Hydrogen-deuterium exchange mass spectrometry (HDX-MS) has become an exceptional tool for probing structure and dynamics of proteins, nucleic acids, and other chemical systems. A general workflow of a ‘bottom-up’ HDX-MS experiment involves the exchange of a protonated sample in a deuterated buffer for different exposure times ranging from milliseconds to hours followed by rapid quenching, proteolysis, and liquid chromatography mass spectrometry (LC-MS) analysis (Masson, Burke et al. 2019). Other experimental setups are possible (Mistarz, Bellina et al. 2018, Wollenberg, Pengelley et al. 2020), but the key output is a mass spectrum in the form of mass over charge (m/z) versus intensity for each analyte (typically peptides), modification state (or general chemical formula), charge state, and deuterium exposure time. The spectra will typically follow a binomial distribution based on the underlying natural abundance isotopic envelope and the level of deuterium uptake (Chik, Vande Graaf et al. 2006).

For native protein samples a given spectrum may reflect a single underlying deuterium exchange state and be sufficiently fit by a single (unimodal) binomial distribution, but it is also possible for multiple exchange states to be present resulting in a multimodal binomial distribution. Multimodal analysis of HDX-MS data is not routine and is often only implemented in cases where spectra show clear separation between exchange states (e.g. “pass the eye test”, Figure 1A bimodal and trimodal spectra). Analysis of multimodal data has relied on a variety of existing software or simple Gaussian fits (Weis, Engen et al. 2006, Pascal, Willis et al. 2012, Guttman, Weis et al. 2013, Kan, Ye et al. 2019, Na, Lee et al. 2019, Largy and Ranz 2023) and is generally applied only to select spectra and can be very time consuming. A key challenge of applying this analysis to a whole dataset is determining when to fit each spectrum with multiple populations and the subsequent extraction of the fit parameters, chiefly the respective deuterium uptake values and the populations of the underlying states.

**Figure 1.**
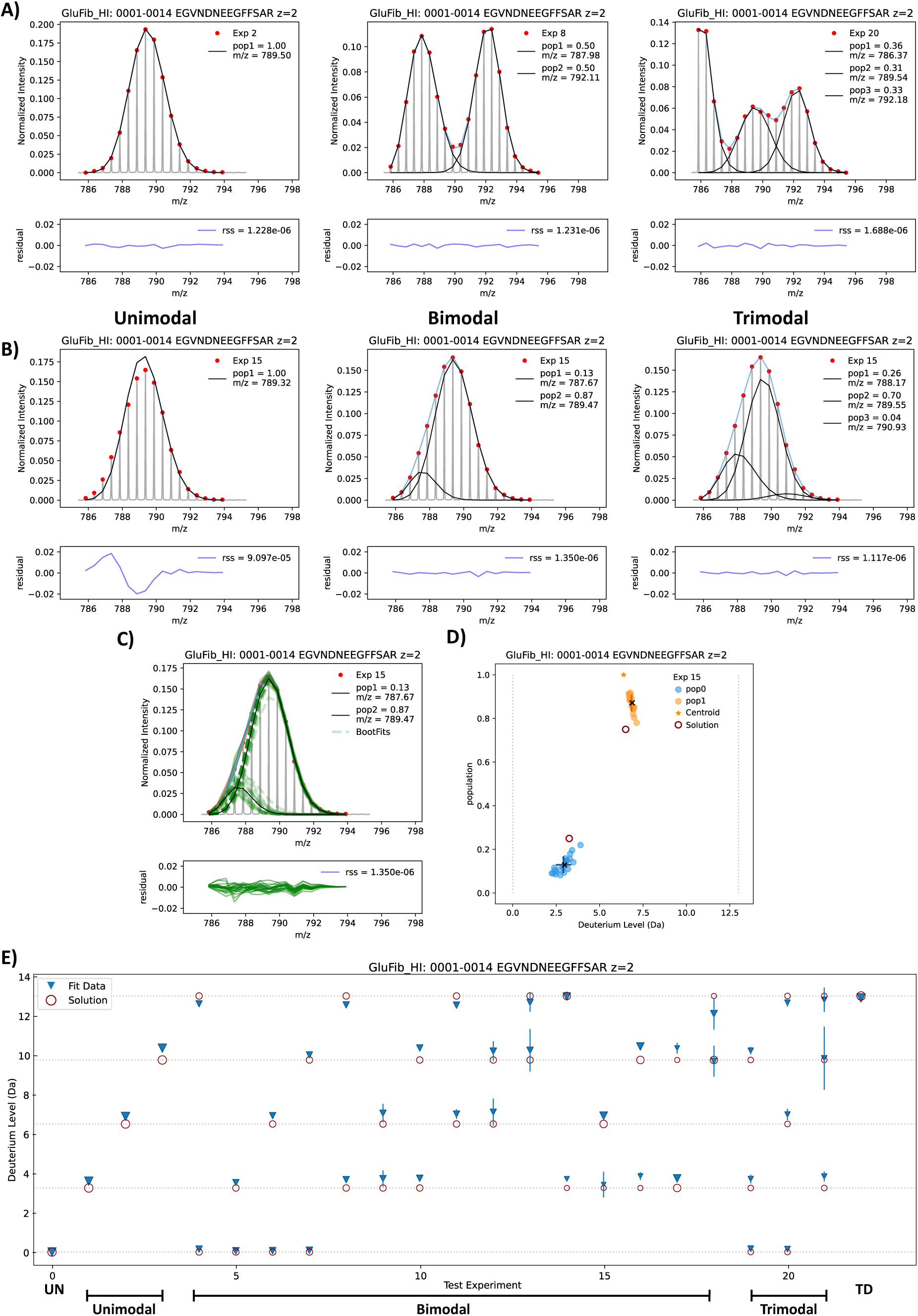
A) Known mixtures data with one, two, or three components, with clear separation. The raw data is shown in grey, picked peaks are red circles, the total fit is a blue line, and the underlying unimodal, bimodal, or trimodal fits are shown as black lines. The lower plots represent the residual error at each data point and indicate good agreement between the fits and experimental data. B) A 1:3 mixture of 25% and 50% deuterated samples. The raw data appears unimodal to the eye but is fit poorly by a single population. Bimodal or trimodal fits improve the RSS but the improvement for the trimodal fit (p_fit_ = 0.44) is insufficient to justify fitting additional parameters, so the bimodal fit (p_fit_ = 1.2e-10) is selected. C) The data with added 5% random error is fitted 20 times (BootFits, green lines) and the resulting populations and Deuterium uptake is plotted (right). Right: Red circles indicate the known mixture expected values, which are closely approximated by the bimodal fits. D) Autonomous fits of the full set of known mixtures data for GluFib, allowing up to 4 populations, correctly select the number of components. The number of fit populations is selected based on p_fit_ < 0.05 and pop_min_ > 0.03. Mean and standard error of the uptake values are from 20 BootFits as in C). ‘UN’ and ‘TD’ denote the undeuterated and fully-deuterated controls. Plots were generated as outputs of the pyHXExpress fitting process.

We recently released HX-Express v3 which is a Microsoft Excel-based tool that has established algorithms and statistical metrics for evaluating the multimodality of HDX-MS spectra (Weis, Engen et al. 2006, Guttman, Weis et al. 2013, Tuttle, James et al. 2025). The HX-Express implementation is applied to spectral data one-by-one on a single analyte in a semi-automated manner. We adapt this approach to implement a high-throughput and fully-automatic multimodal analysis of HDX-MS spectra in a freely available Python package called pyHXExpress (https://github.com/tuttlelm/pyHXExpress). Here, we show that pyHXExpress can autonomously and correctly fit the number of underlying states and their deuterium uptake levels and populations of a previously reported test dataset of known mixtures (Tuttle, James et al. 2025).

Previous multimodal analysis of deuterium exchange spectra has typically involved only a handful of peptides in each system (Clouser, Baughman et al. 2019, Costello, Shoemaker et al. 2022, Calvaresi, Wrobel et al. 2023, Torres-Paris, Song et al. 2024), and there is not an established protocol for handling a multimodal analysis of the massive amount of data contained in a full protein dataset. To meet this need for whole system bimodal characterization, we propose a new statistical approach that leverages all the spectra that contribute to the deuterium exchange information of each residue. As a demonstration of this approach, we use pyHXExpress to analyze more than 5,000 spectra and characterize the bimodality in the entire systems for the small heat shock protein HSPB5 and two disease mutants D109H and R120G. From this we gain new insights into the pervasive bimodality throughout the N-terminal region (NTR) of wildtype (WT) HSBP5 and the mutants, are able to directly compare these species, and gain insights that are not revealed from a unimodal analysis (Woods, Janowska et al. 2024).

## METHODS

Full guidance on installation and use of pyHXExpress is available at https://github.com/tuttlelm/pyHXExpress. An overview of the analysis workflow and fitting logic of pyHXExpress is given as Figure 2 and is described here.

**Figure 2.**
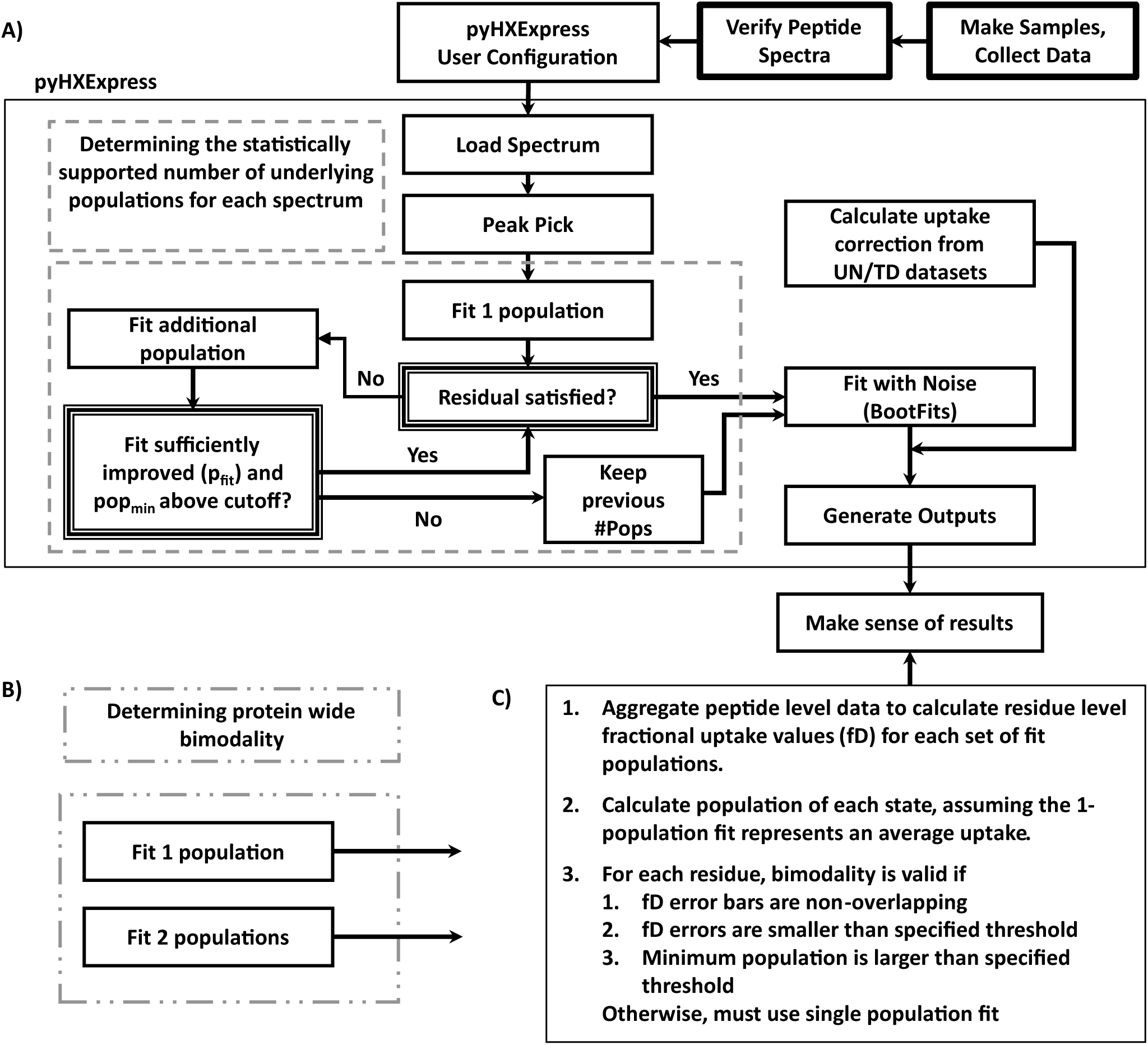
Flow diagram of the HDX-MS data fitting procedure using pyHXExpress. A) The overall scheme is shown, with the specific protocol for choosing the correct number of fits in a single spectrum indicated by the dashed boxes. For this approach, the user specifies the minimum and maximum number of underlying populations for each fit, a p_fit_ cutoff, a minimum population threshold, and the RSS cutoff, which are used to evaluate whether additional fit populations are justified. To evaluate the protein wide bimodality, the dashed boxes of B) are substituted for those in A). All datasets are fit to 1-population and to 2-populations and are evaluated according to C) to determine the bimodality at the residue level (minimum resolvable peptide block). This analysis should be performed in combination with the data quality metrics resolution (size of the minimum peptide block contributing to the residue level fD) and redundancy (the number of spectra contributing to the residue level fD).

### Input Data

At the most basic level, pyHXExpress takes as input raw data of m/z versus intensity for each sample/peptide/charge/timepoint/replicate. This can be in the tabular format recognized by HX-Express or, conveniently, the entire peptide pool exported spectra from HDExaminer (Trajan Scientific and Medical). A metadata dataframe (metadf) .csv file is used to control what data will be fit. It can be automatically generated if using the HDExaminer outputs and additional FASTA files for each species but otherwise it must be created by the user. This file specifies, for example, the peptide sequence, charge state, and peptide modifications for each data file. The modifications designator is particularly powerful as this allows pyHXExpress to be used with HDX-MS data from any system, not just proteins, as long as the correct chemical formula and number of exchangeable protons (Hex) for each spectrum is specified. The last requirement is a user configuration file that provides file path information and a variety of parameters that control how the fits should be performed, including e.g. the minimum and maximum number of populations to fit, a p-value cutoff to accept additionally fit populations, and the residual sum square cutoff indicating when a fit should be considered good enough.

### What pyHXExpress does

The general procedure of pyHXExpress (Figure 2A) is to 1) read in the spectra, 2) pick the peaks at the expected m/z values (Kan, Ye et al. 2019), 3) fit the data to the non-integer (real numbers) version of the binomial distribution using the underlying pattern of the natural abundance isotopic mass envelope (Dittwald and Valkenborg 2014) using the user specified number of initial guesses (‘BestFit_of_X’) and take the fit with the lowest residuals, 4) evaluate the quality of those fits, which determines whether an additional population is fit based on user settings, and repeat, and 5) with the number of underlying populations determined, perform a resampling method whereby the data plus some amount of random noise is refit a specified number of times (‘Nboot’, default 20). This resampling, which is implemented in the most recent version of HX-Express (Tuttle, James et al. 2025), is a regularization method to reduce overfitting and allows for calculation of meaningful errors on each fit parameter. We term these resampled fits ‘BootFits’, as the method is inspired by the statistical Bootstrap method. The size of the noise added to each point is drawn from a Gaussian distribution with the maximum value specified by the user (either as an absolute intensity value or as a percentage of the maximum intensity).

When multiple populations are fit, by definition the underlying fits are grouped from slowest exchange to fastest exchange (so pop1 will be the population with the slowest exchange, pop2 next slowest, etc.). For each sample/peptide/charge set of spectra, the back-exchange correction is calculated from the unimodal fits of the undeuterated and fully-deuterated control spectra (Masson, Burke et al. 2019). It is possible to perform fits without these controls, though the outputs will be flagged accordingly: the centroid of the natural abundance isotopic envelope substitutes for the undeuterated control and the user-specified fraction of deuteration in the exchange buffer times the expected number of exchangeable amides substitutes for the fully-deuterated control. Available outputs include but are not limited to figures and tables of the fits, and single file outputs of the raw data and peak-picked data.

### Performing the fits

With the required input files in place, the user can run pyHXExpress scripts to fit the HDX-MS spectra and generate the output plots and tables of fit parameters. As a single protein can have thousands of spectra, this process is expedited by performing batch fits where subsets of the metadf-specified datasets are processed in parallel. For reference, the complete fits of more than 5,000 spectra for the HSPB5 WT, D109H, and R120G mutants, for both fixed 1- or 2- population fits, took roughly 24 hours when batched as 6 separate jobs (AMD Ryzen Threadripper 3960 processor). The minimum batchable unit is all timepoints and replicates of a single peptide and charge state, since back-exchange corrections are applied based on the corresponding undeuterated and fully-deuterated control samples; this is generally what is specified by a single line entry of the metadf file.

### Validation against test data

We verify the implementation of pyHXExpress by performing fits on known deuterium uptake mixtures of the glufibrino peptide (GluFib; peptide sequence EGVNDNEEGFFSAR; charge +2) as described previously (Tuttle, James et al. 2025). This dataset includes 21 different mixtures of known deuterium uptake (Exp 1-21) plus the undeuterated (Exp 0) and fully-deuterated (Exp 22) control spectra. Each spectrum was fit with 1 to 4 populations, and the correct fit was selected based on the criteria of Figure 2A, using p_fit_ < 0.05 and min_pop_ > 0.03. The residual cutoff was set to 0, so it was not used in fit selection.

### Full protein bimodal analysis

Instead of evaluating bimodality at the single spectrum level, all available spectra for a given residue inform on whether bimodality is supported for that residue (Figure 2B-C). The back-exchange corrected relative fractional D-uptake (fD) is calculated for each spectrum. pyHXExpress outputs can be used with other HDX-MS analysis suites such as HDXBoxeR (Janowska, Reiter et al. 2024) and pyHDX (Smit, Krishnamurthy et al. 2021). We utilize the residue-level D-uptake calculation method of pyHDX using a forked version that has been adapted to allow multiple replicates (https://github.com/tuttlelm/PyHDX). This results in “residue level” D-uptake averages and standard deviations for each exposure time. The actual residue level resolution depends on the peptide coverage, where the minimum resolved peptide block is a function of peptide overlap. We calculate the fD_avg_ value from the fixed 1-population fits and fD_fast_ and fD_slow_ from the fixed 2-population fits. For this analysis we make the following assumptions which will need to be evaluated on a case-by-case basis: 1) the 1-population fit is an appropriate proxy for the average exchange and 2) all slower exchange populations correspond to a slow state of the protein while all faster exchange populations correspond to the fast state. Assumption 1 is reasonable so long as the fit is not clearly missing a well-resolved subpopulation (as may occur if a single population is fit to the Figure 1A multimodal examples). A simple alternative would be to use the centroid as the average proxy, though this will be more susceptible to contaminating peaks. Assumption 2 is expected to be generally reasonable for residues close in sequence space, however with external data about the system, the state 1 (here, all slow) or state 2 (here, all fast) assignments could be adjusted on a per peptide basis manually. Although the fits include the populations as a fit parameter, these values are inherently more error prone and vary widely from peptide to peptide as whatever underlying exchange processes between substates may be occurring will differently affect the observable populations. Instead, we use the relationship between fD_slow_, fD_fast_, and fD_avg_ to derive a surrogate population measure, such that

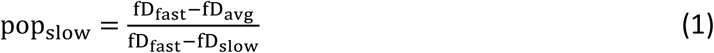

and pop_slow_ + pop_fast_ = 1. Errors for fD and pop values were propagated from underlying values using the propagation of variance approach:

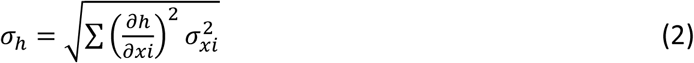

where *h* is a function of the components *x_i_*. For each residue, the requirements to accept the bimodal fit as appropriate are the following: 1) fD_fast_ and fD_slow_ should fall between 0 and 1, 2) fD_avg_ must fall between the values of fD_fast_ and fD_slow_ 3) the error ranges of fD_fast_ and fD_slow_ should not be overlapping, 4) the error in either fD value should be less than some cutoff (e.g. 0.3), and 5) the derived minimum population for one state should be larger than some value (e.g. 5%). If all criteria are not met, a bimodal fit cannot be supported by the data and the unimodal 1-population ‘average’ fit is selected.

These data are qualitatively assessed based on the resolution (number of residues in the minimal residue block) and redundancy (number of measurements contributing to the values) for each residue. Resolution is largely dependent on peptide cleavage which determines the peptide coverage and peptide overlap. The redundancy depends both on peptide overlap (i.e., one residue occurring in multiple peptides) and the number of replicates collected.

This approach is applied to the recently published HDX-MS data of the small heat shock protein HSPB5 and mutants D109H and R120G at pH 7.5 where there are significant differences in activity between WT and the mutants (Woods, Janowska et al. 2024), to ascertain whether bimodality is statistically supported at each residue. The data were collected with four replicates at 4, 60, 1800, and 72000 sec exposure times plus the undeuterated and fully-deuterated control samples.

## RESULTS AND DISCUSSION

### Autonomous fitting of multimodal HDX-MS data

pyHXExpress is a Python implementation of the fitting subroutines used in HX-Express v3 with additional functionality that allows high-throughput fully automated analysis of complete sets of HDX-MS spectra. The workflow is as shown in Figure 2, and we validate this approach against an existing test set of peptides with one to three components of known deuterium exchange (Tuttle, James et al. 2025). In some cases, as in Figure 1A for the GluFib peptide in various mixtures, the agreement between the fit and the data is readily discernable, even for bimodal and trimodal spectra. This is generally not the case in real data where distinction among potentially different states is less clear. Additionally, thousands of spectra will potentially need to be fit and evaluated with a robust and quantitative statistical approach. We have recently established a primary evaluation metric using p_fit_ (p-value from the F-test) which evaluates whether fitting an additional population is statistically supported based on the improvement in the residual sum of squares (RSS), with a typical requirement of p_fit_ < 0.05. For the F-test to be valid, it is essential that the compared fits be the best achievable fit (global minimum of RSS) for that number of populations. To achieve this, fits are performed multiple times (user-specified ‘BestFit_of_X’) from different initial conditions and the parameters giving the lowest RSS are selected. We find that using a BestFit_of_X of at least 3 is required to choose the correct number of underlying populations, with diminishing improvements as this value is increased.

The improvement to the fits as we allow more parameters (populations) to be fit is shown in Figure 1B. This spectrum is for the GluFib peptide in a 1:3 mixture of 25% and 50% deuterated components. The picked peaks may appear unimodal to eye, but the 1-population fit poorly reproduces the early data points, which is reflected in a large residual for those points. The bimodal fit significantly improves agreement with the experimental data with a p_fit_ of 1.2e-10. Fitting three populations gives modest improvements to the RSS but comparison to the bimodal fit gives a p_fit_ of only 0.44, indicating that the additional parameters are not statistically justified. To reduce computation time, pyHXExpress has additional user-defined criteria for choosing the appropriate number of populations to fit, which include an RSS threshold for “good enough” and a minimum population requirement. Both measures can also curtail fitting of spurious contaminating peaks. For this small dataset of high quality GluFib peptide data, no residual cutoff was used (set to 0), while the pop_min_ > 3% was sufficient to avoid overfitting noise peaks.

With the number of underlying populations determined, we perform a data regularization step inspired by Bootstrap resampling (that we term ‘BootFits’) whereby some amount of random noise is added to the experimental data and refit 20 times (specified by user). This approach also allows for generation of meaningful error ranges determined from the standard deviation of these BootFits. The BootFits are shown as the green curves in Figure 1C, with clear clusters of fits for the less populated less exchanged state and for the more populated more exchanged state. The population versus deuterium uptake plot in Figure 1D shows the fit parameters versus the expected ‘solution’ (which will also have error bars of unknown size). The fits very closely approximate the expected components of a 25% population of 25% deuteration and a 75% population of 50% deuteration. The complete GluFib test set includes 21 different mixtures of known deuteration state (Exp 1-21) and the undeuterated (UN, Exp 0) and fully-deuterated controls (TD, Exp 22). The uptake values are corrected such that the undeuterated sample has 0 D-uptake and the fully-deuterated sample has 13 D-uptake (14 non-proline residues less 1 N-terminal residue). When pyHXExpress is run on this dataset according to the criteria of Figure 2A using the p_fit_ < 0.05 and pop_min_ > 0.03, we correctly identify the number of underlying components and their D-uptake levels for all 21 mixtures (Figure 1E). This demonstrates that when the data are high-quality it is possible to correctly characterize the underlying deuterium exchange states.

It is important to note that when fitting real data (which will have both random and systematic error), this approach answers the question of whether it is statistically justified to assume additional populations based on a single spectrum, which is different than absolutely determining the underlying exchange states. The case may be that there are additional populations, but the data are of insufficient quality to quantitatively identify them. It is also especially important to have high quality spectra with minimal peptide contamination. Because the purpose of this analysis is to detect multimodal data, it is necessarily assumed that any intensities at the expected m/z values are real. pyHXExpress outputs will flag cases where the undeuterated or fully-deuterated controls appear multimodal, which can be a sign of contaminating peptide peaks.

### Full protein bimodal analysis of HSPB5

We next show how pyHXExpress enables the evaluation of bimodal character throughout a protein. One approach would be to evaluate the number of underlying populations for every spectrum as was done for the GluFib test set (Figures 1 and 2A). This is a reasonable way to assess overall data quality and can highlight regions where multimodal fits are statistically supported. What quickly becomes apparent, however, is that different replicates or charge states of a single peptide and exposure time might support a different number of underlying populations. This can be due to the sample-to-sample variability for different replicates or due to differences in signal to noise or peptide overlap for different charge states. As comprehensively computing the multimodal fits of all peptides has not been feasible before now, there is no established protocol for determining the statistically appropriate number of underlying populations. Current metrics, as described for the GluFib peptide, rely on statistical tests against a single spectrum; however, multiple peptides, charge states, and replicates can report on the exchange properties of each residue. As such, we propose a method that aggregates all peptide spectral data to determine whether multiple populations are supported at each residue (Figure 2B-C).

We present this approach applied to the small heat shock protein HSBP5 as a test case. HSPB5 is a chaperone protein expressed in many tissue types including eye lens, skeletal and cardiac muscle, and brain. Inherited missense mutations, D109H and R120G, are associated with similar pathologies including myopathies and early cataract. HSPB5 is comprised of a quasi-ordered N-terminal region (NTR, residues 1-63), a central structured ɑ-crystallin domain (ACD, residues 64-148), and a disordered C-terminal region (CTR, residues 149-175). Importantly, HSPB5, like other small heat shock proteins in the human proteome, form large, heterogenous, polydisperse oligomers, rendering them intractable by conventional structural biology approaches. It has been established that the disordered NTR is required for oligomeric assembly and for chaperone activity (Jehle, van Rossum et al. 2009).

The two disease mutations investigated both reside within the structured ACD, yet they display differing chaperone activity, begging the question of how the mutations affect the NTRs within oligomers (Woods, Janowska et al. 2024). The conventional centroid analysis of the HDX-MS spectra of WT HSPB5 and two mutants was highly informative, revealing differences among the three proteins (Woods, Janowska et al. 2024). The known heterogeneity of NTRs within HSPB5 oligomers (Jehle, Vollmar et al. 2011) makes the protein and its disease mutants an interesting system in which to detect and compare underlying exchange states. HSPB5 likely contains numerous underlying exchange states, but we find that generally just two states can reliably be resolved and considered in any consistent way throughout the protein sequence.

It is important to be aware of factors that may lead to detecting more than one population. The first consideration is quality of the underlying spectra. They should have signals reasonably above the noise, with peaks at the expected m/z values, and no contaminating peptides. Admittedly, this is non-trivial to achieve for the thousands of spectra obtained for even a single protein state. In an ideal case, we would have several replicates of each experiment that give high quality spectra and have variable length peptides that cover the whole sequence, with massive overlap differing by only one residue for each successive peptide. In reality, this is not the case, and while we have reasonable control over the number of replicates collected (within the limits of sample, personnel, and instrument time), we have far less control over the peptides generated for each system, although this can be optimized by using different or multiple proteases. We rely on the residue level redundancy and resolution measures to assess the quality of the ‘residue level’ derived values (Figure 3B) (Smit, Krishnamurthy et al. 2021, Stofella, Grimaldi et al. 2024). Redundancy is the number of spectra that contribute to the measurement of the fractional D-uptake (fD) including replicates, and resolution is the size of the minimum peptide block that is used to derive these values. The fD versus exposure time uptake plots are shown for each of these minimal blocks in Figure S2. For HSPB5, the lowest resolution is in regions within the ACD and CTR, visible as red bars in the resolution plot of Figure 3B. The peptide coverage, resolution, and redundancy are very similar for HSPB5 WT, D109H, and R120G which facilitates comparison of results among these species.

**Figure 3.**
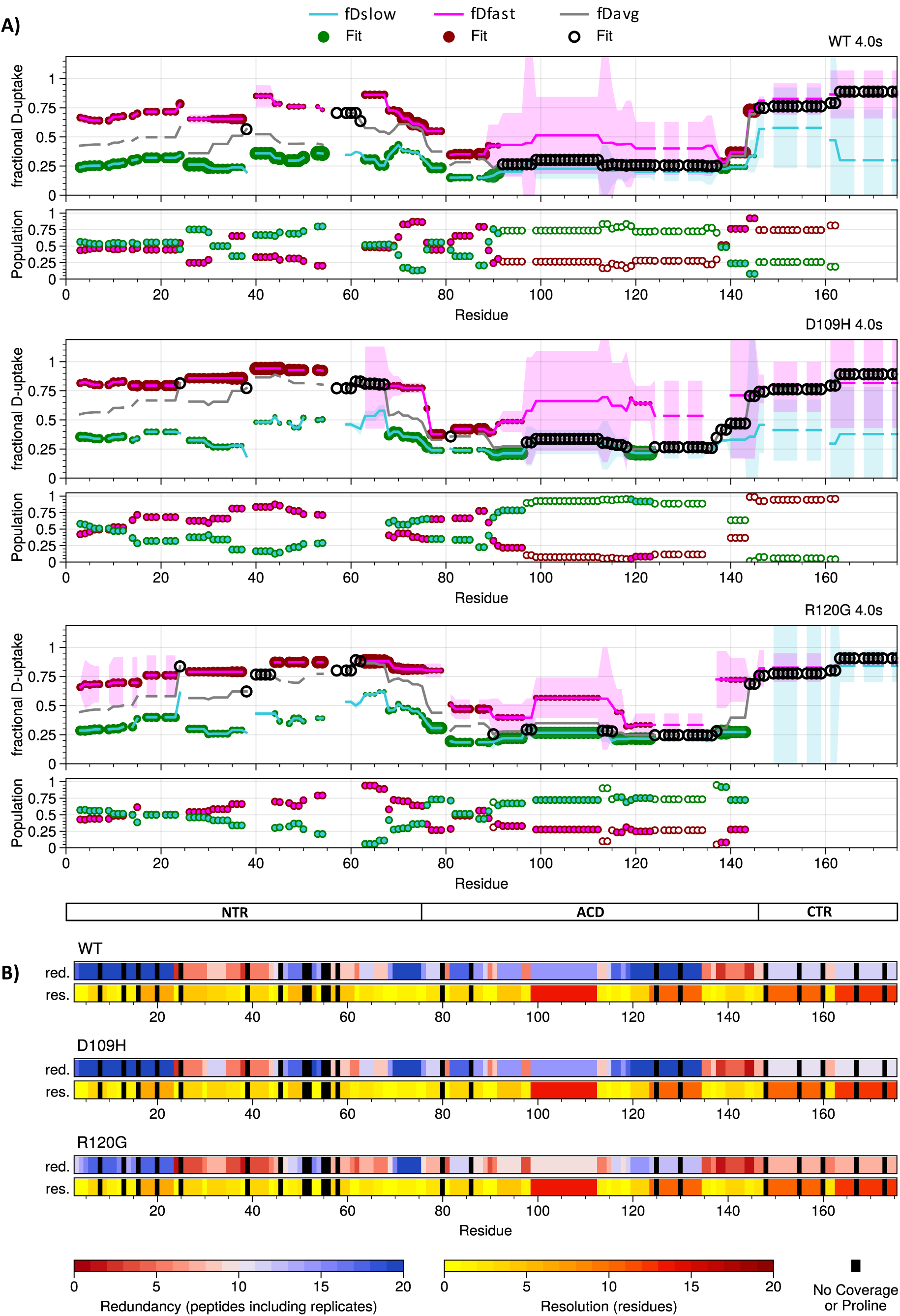
A) Comparison of the fractional D-uptake and populations for the 1-population fit (fD_avg_) and 2-population fits (fD_slow_ and fD_fast_) for HSPB5 WT, D109H, and R120G at the 4 sec exposure timepoint. If the bimodal fit is selected (criteria in Figure 2C) a filled circle is shown for that residue where the size scales with population, otherwise an open black circle denotes where the unimodal fit is selected. Standard deviations of the residue uptake values are shown by shaded areas. In the population plots, open circles denote regions where the 1-population fit is selected. The HSPB5 domain architecture is shown at bottom, with subregions of the NTR separated by dashed vertical lines. For each species, 2- populations are supported for much of the NTR, and a single population is supported for the CTR. B) The redundancy and resolution for each species. This applies to all exposure times since only peptides with data at all exposure timepoints were retained. A higher redundancy means more spectra contributed to the calculations of D-uptake and a lower resolution means that value represents a smaller minimal peptide block. Black bars indicate regions that are proline or have no peptide coverage.

HSPB5 presents a case where there are clear regions of multiple exchange states and other regions where only a single population fit is justified from the data. In Figure 3A, we show the comparison of the 2-population fits (fD_slow_ and fD_fast_) and the 1-population fit (fD_avg_) for each HSPB5 species for the 4 sec deuterium exposure time. At each residue, a circle indicates the fit that has been selected based on the criteria of Figure 2C. The plots of Figure 4 are meant to facilitate comparison of the deuterium exchange of each HSPB5 species. Figure 4A shows an overlay of the fractional D-uptake results for the 4 sec exposure time, with only data of the appropriately selected fit shown (fD_fast_ and fD_slow_ are lines, fD_avg_ are circles) along with the population of the more protected state corresponding to fD_slow_. The plots make clear that, in general, the fD_slow_ and fD_fast_ levels are similar among the three species, but the distribution between the slow and fast exchanging states differ in the mutants. In particular, the more protected state of residues ∼15-30 is more lowly populated in D109H than either WT or R120G, while both mutants de-populate the protected state for residues ∼40-55. These observations provide invaluable insights regarding the mechanism of HSPB5 and the consequences of the mutations, but this is outside the scope of this paper.

**Figure 4.**
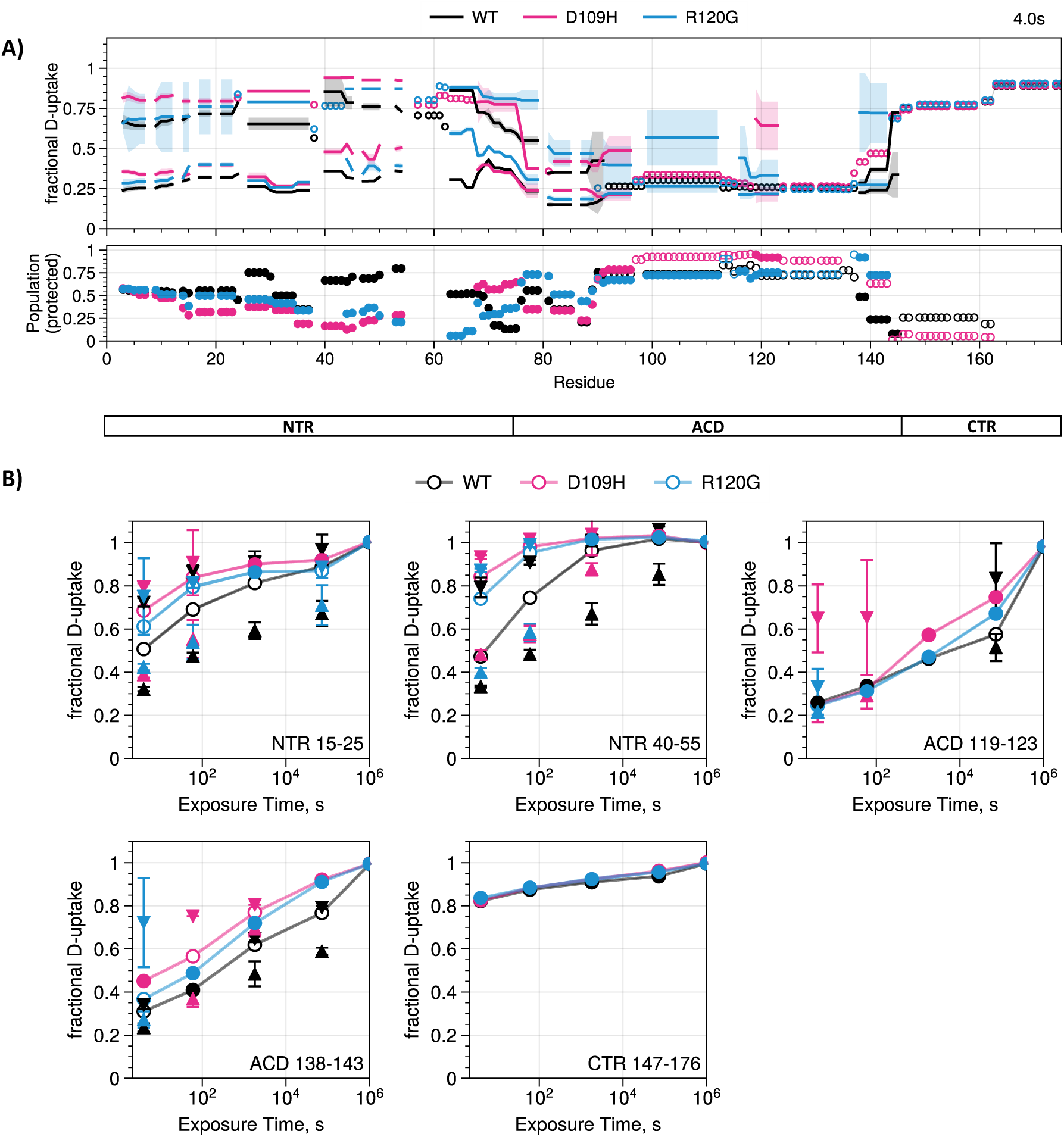
A) Overlay of the fractional D-uptake and population plots for WT (black), D109H (red), and R120G (blue) at the 4 sec exposure timepoint. For each residue, lines are shown if the 2-population fits were justified or as open circles if the 1-population fit is selected. The population plot shows the population of the more protected state corresponding to fD_slow_; open circles are shown if the 1- population fit was selected. The NTR of all species shows widespread bimodal behavior with similar underlying exchange states that are differently populated. The ACD region of all is more protected, but a lowly populated faster exchange population is supported for R120G. For all species, the CTR beyond residue 145 is similarly rapidly exchanged. B) Uptake versus exposure time for select regions throughout HSPB5, a dummy time of 10^6^ sec is used for the fully-deuterated experiment. fD_slow_ is shown as upward triangles, fD_fast_ is downward triangles, and fD_avg_ is shown as lines. Filled triangles indicate that the 2-population fit is supported while filled circles show where 1-population is supported.

The bimodal analysis reveals that much of the NTR has both faster and slower deuterium exchange populations that are detectible throughout the exposure time course (Figures 3, 4, S1, and S2). In contrast, the CTR is rapidly exchanged as a single population even at the shortest 4 sec exposure. Beyond residue 90, the ACD is more protected as a single population in all cases, except for R120G, which has a lowly populated faster exchange population detectible at the shortest exposure time, and for D109H near residue 120 (the ACD dimer interface) where a very lowly populated more exchanged population is also detected. The dimer interface only becomes detectible as two exchange populations for WT at the longest exposure time, which may be reporting on the slow subunit exchange of WT oligomers, as it takes a long time for the more exchanged state to become populated. The region near the ACD to CTR transition also detects bimodal species in a clear example of peptides containing mixed secondary structure elements (the more protected ACD and the disordered CTR).

This data highlights the types of scenarios that may be detected in a system with regions sampling conformations with different levels of solvent protection. When the more protected conformation visits the less protected conformation, deuterium uptake will occur such that the observable population of the protected state decreases. This is evident, for example, for the mutants in regions of the NTR where at early timepoints two exchange populations are detected, but as the exposure time increases, the population of the protected state decreases below detectability and the unimodal fit is selected (Figure S1B,C). On the other hand, as indicated for the WT dimer interface, when a region is predominantly protected, it may take some time for the faster exchange state to grow in population to detectible levels.

A key question in this full protein bimodal analysis is what can we learn that is not revealed by the far simpler and faster unimodal ‘average’ analysis? We address this question here in a more illustrative proof-of-concept way than as an exhaustive analysis of all we can learn about HSPB5 from these data (a not unworthy endeavor, but beyond the scope of this current work). As mentioned above, the NTRs of WT and mutant HSPB5 each have similar protected and more exchanged states, with the greatest differences being in the relative populations of these states (Figure 4). The greatest difference in these populations is for residues ∼15-25 and residues ∼40-55, where WT has a much larger protected population than either mutant. Both these regions are known from other experiments to contact the structured ACDs, binding into grooves presented on the ACD surface. A plausible model is that the more protected exchange state reflects the population of groove-bound sequences within oligomers. Notably, both disease mutations are in one such groove, providing a possible explanation for why these mutations alter the distribution of NTR regions to favor the less protected states. The quasi-ordered NTR is known to have multiple competing interactions throughout HSPB5, and it may be that the mutants have altered the balance of these interactions relative to WT, which tends to be more protected on average. While not conclusive from these data alone, this is consistent with a similar underlying conformational landscape with different exchange rates between these states.

## CONCLUSIONS

We have created a freely available Python codebase, pyHXExpress, that incorporates all the statistical rigor of HX-Express while enabling the high-throughput autonomous multimodal analysis of complete HDX-MS datasets. Currently, multimodal analysis of even select peptides in a protein system is rare, and the comprehensive bimodal analysis of all peptides in a native protein system has not been reported. pyHXExpress is a tool that will enable this to become more routine. As a test case and proof-of-concept, we have applied the full protein bimodal analysis to the small heat shock protein HSPB5 and two disease mutants D109H and R120G. To do this, we present a new statistical approach to assess bimodality that leverages all peptides and charge states that a residue occurs in to assess the bimodality at the residue level, with the size of the minimum peptide blocks dictating this residue level resolution. This approach has allowed us to gain new insights into the conformational and dynamic landscape of HSPB5, an infamously challenging protein target for rigorous, residue-level study. We hope that with full protein bimodal analysis now more accessible that this will push the field to address the challenges and opportunities of better understanding the wealth of information contained in HDX-MS datasets.

## AUTHOR CONTRIBUTIONS

L.M.T., R.E.K., and M.G. designed the research. L.M.T performed the experiments and wrote the Python code. L.M.T., R.E.K., and M.G. analyzed data. L.M.T., R.E.K., and M.G. wrote the paper.

## ACKNOWLEDGEMENTS

We thank Damien Wilburn for early coding assistance; Ellie James and Natalie Stone for beta testing; Maria Janowska, Natalie Stone, and Christopher Woods for sharing experimental data to test the code; and attendees of the HDXMS2024 conference for supportive discussions. This work was funded by National Institutes of Health grants: S10OD030237 (M.G.), the National Eye Institute grant 5R01EY017370 (L.M.T., R.E.K.), as well as the National Science Foundation Award: 2304707 (M.G.)

## DATA AND SOFTWARE AVAILABILITY

pyHXExpress is freely available at https://github.com/tuttlelm/pyHXExpress under a GPL-3.0 license. This ‘paper’ branch of this repository includes all Python scripts and Jupyter Notebook files used to generate the analyses and plots presented here.

## SUPPORTING INFORMATION

Supporting information includes Figures S1 and S2. The complete pyHXExpress outputs generated for GluFib and HSPB5 WT, D109H, and R120G are available on the ‘paper’ branch of the pyHXExpress https://github.com/tuttlelm/pyHXExpress/tree/paper.

## SUPPLEMENTARY FIGURES

**Supplemental Figure 1.**
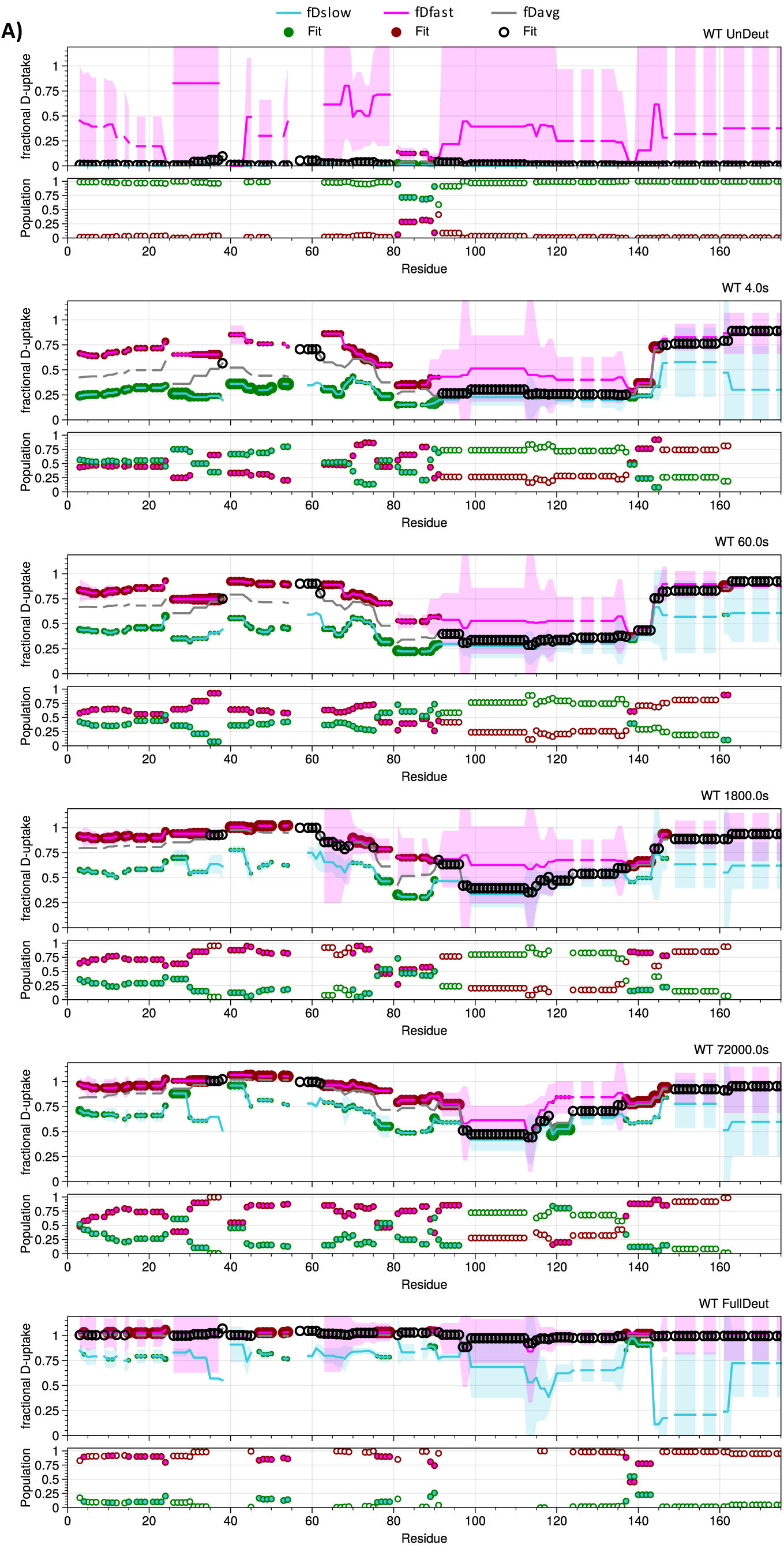
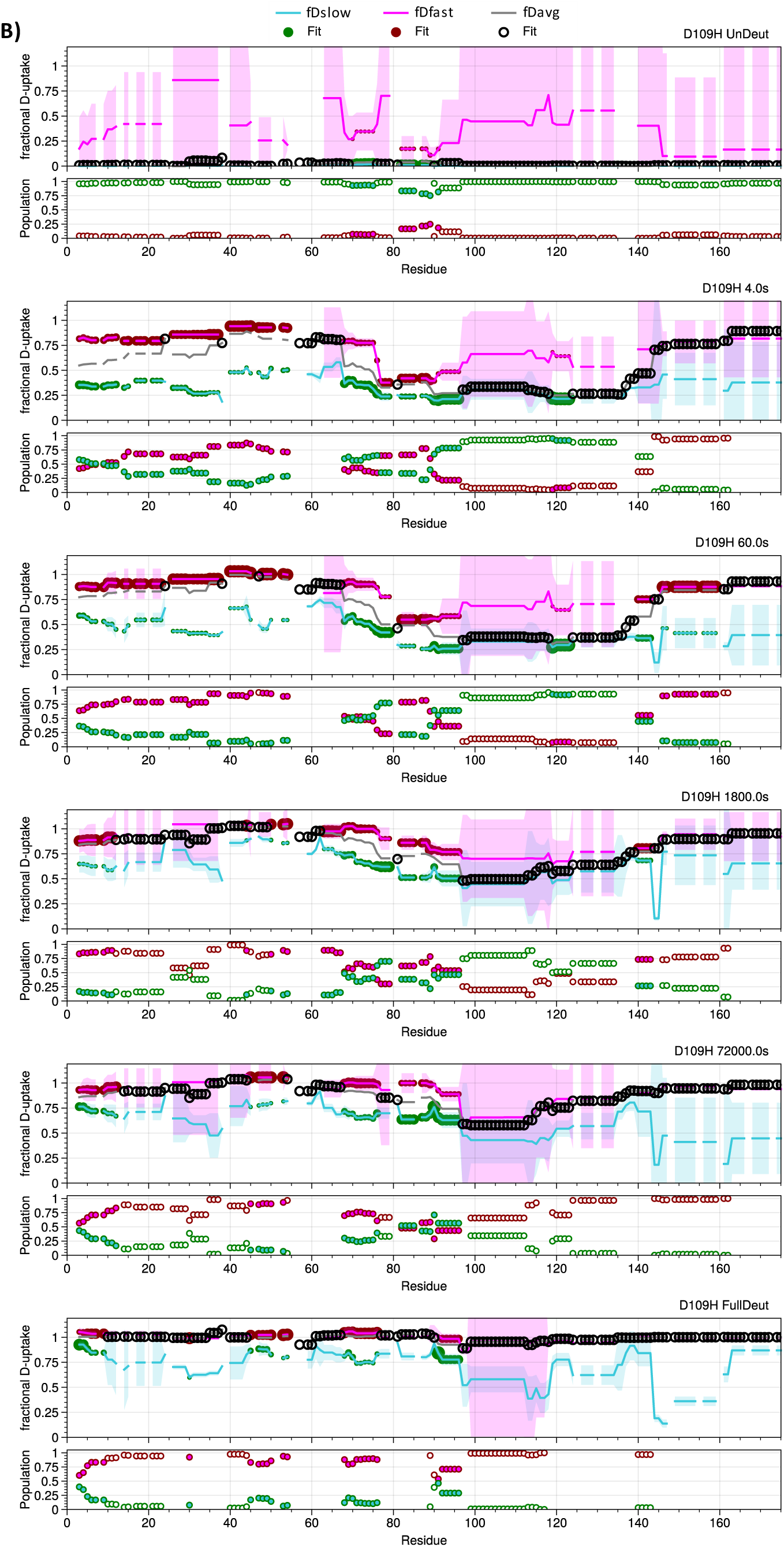
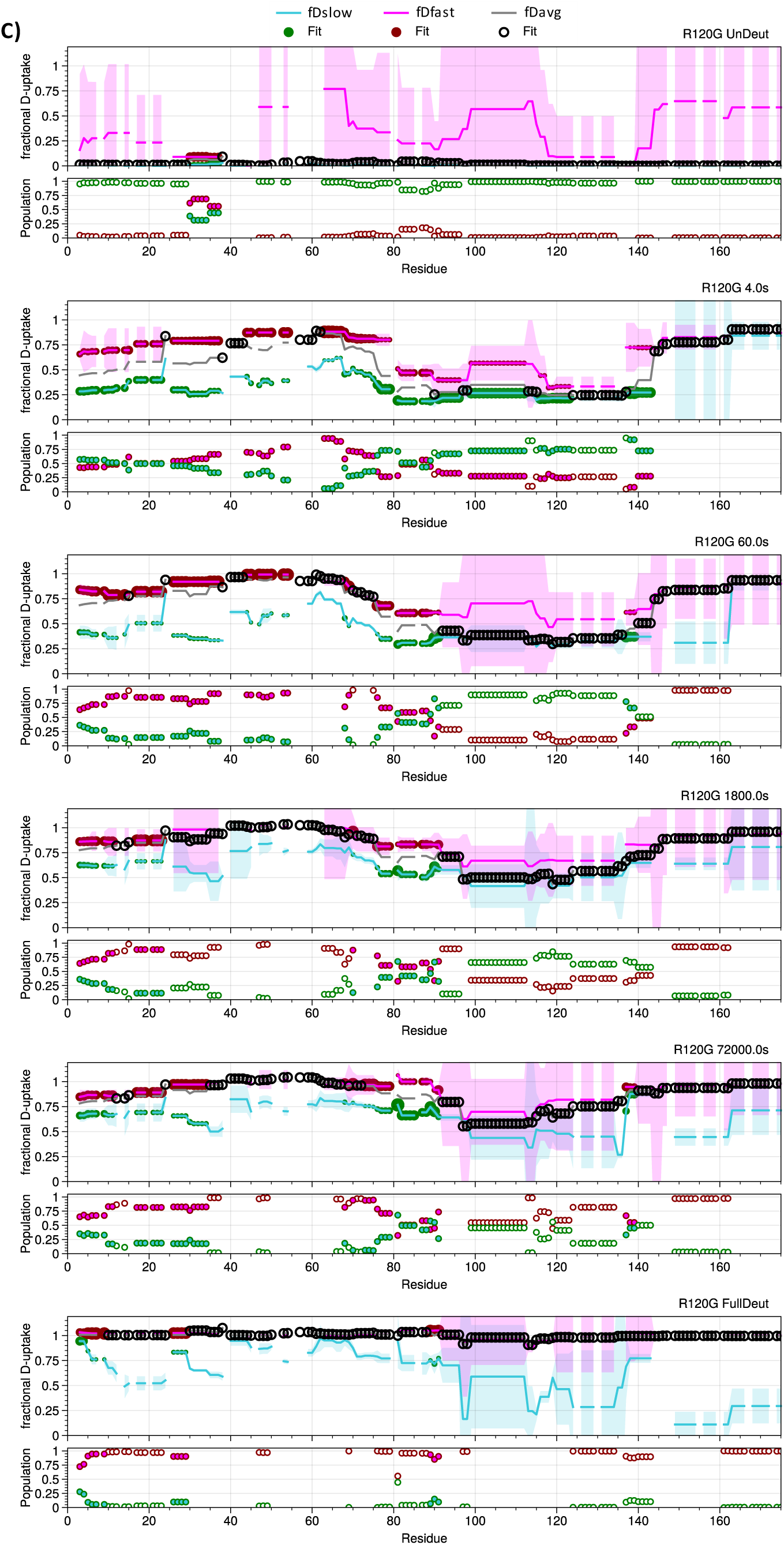
The fractional D-uptake and populations for the 1-population fit (fD_avg_) and 2- population fits (fD_slow_ and fD_fast_) for A) HSPB5 WT, B) D109H, and C) R120G for all exposure times and the undeuterated and fully-deuterated control samples. If the bimodal fit is selected (criteria in Figure 2C) a solid circle is shown for that residue where the size scales with population, otherwise an open black circle denotes where the unimodal fit is selected. Standard deviations of the residue uptake values are shown by shaded areas. In the population plots, open circles denote regions where the 1- population fit is selected.

**Supplemental Figure 2.**
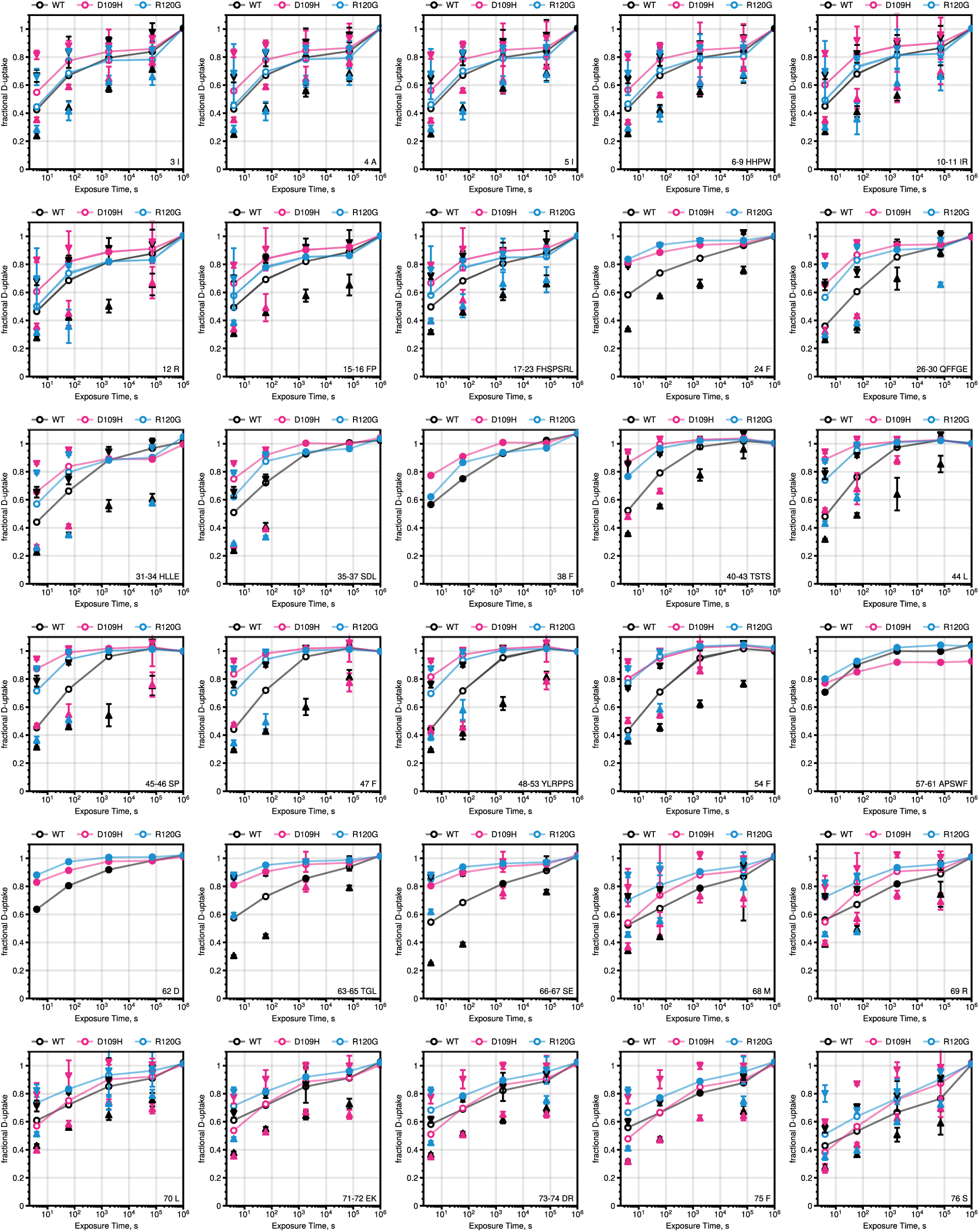
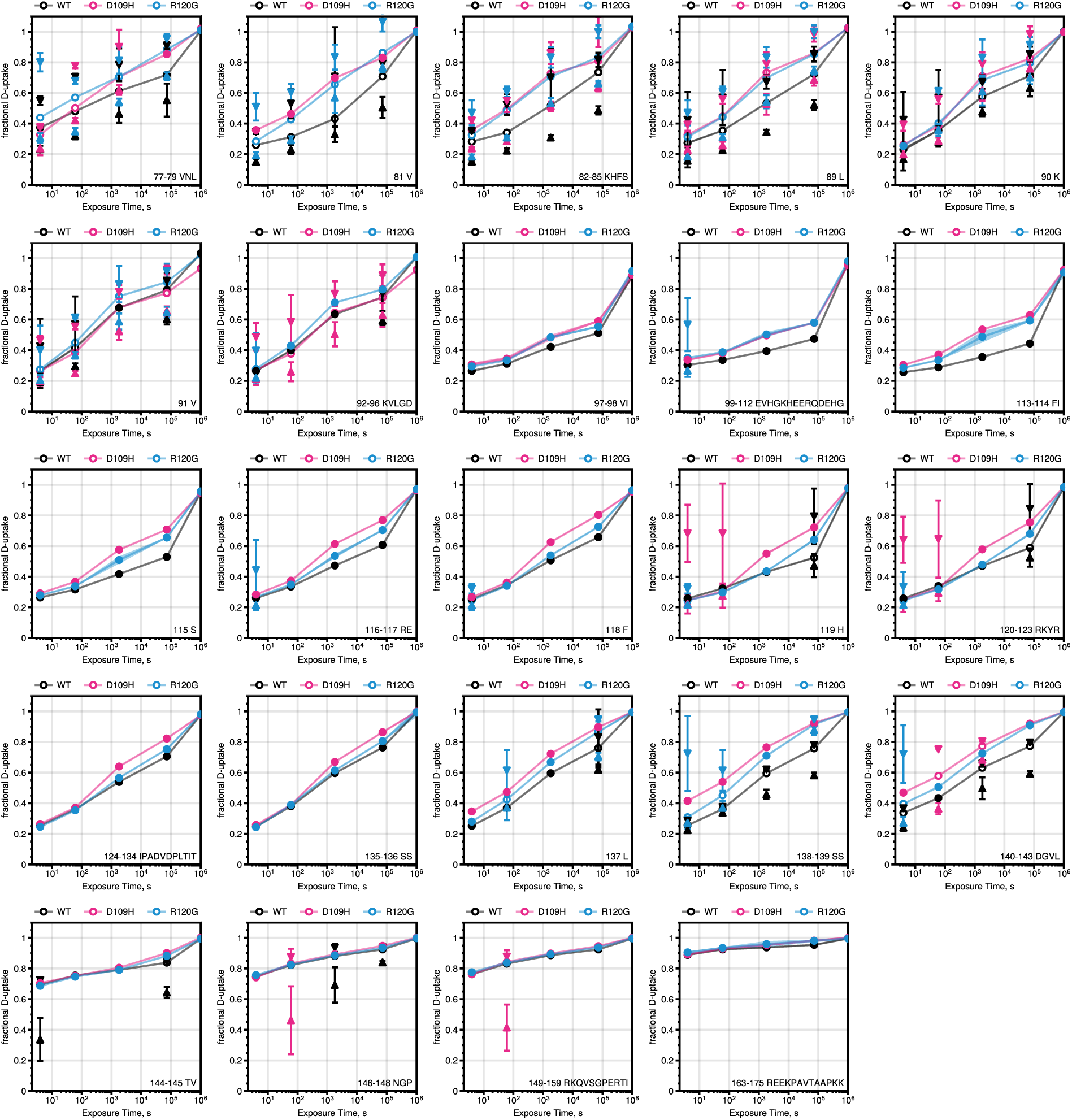
Uptake versus exposure time for each minimal peptide block; a dummy time of 10^6^ sec is used for the fully-deuterated experiment. fD_slow_ is shown as upward triangles, fD_fast_ is downward triangles, and fD_avg_ is shown as lines. Filled triangles indicate that the 2-population fit is supported while filled circles show where 1-population is supported.

## Notes

https://github.com/tuttlelm/pyHXExpress/tree/paper

### Competing Interest Statement

The authors have declared no competing interest.

